# The RNA-binding site of poliovirus 3C protein doubles as a phosphoinositide-binding domain

**DOI:** 10.1101/172742

**Authors:** Djoshkun Shengjuler, Yan Mei Chan, Simou Sun, Ibrahim M. Moustafa, Zhen-Lu Li, David W. Gohara, Matthias Buck, Paul S. Cremer, David D. Boehr, Craig E. Cameron

## Abstract

Some viruses use phosphatidylinositol phosphate (PIP) to mark membranes used for genome replication or virion assembly. PIP-binding motifs of cellular proteins do not exist in viral proteins. Molecular-docking simulations revealed a putative site of PIP binding to poliovirus (PV) 3C protein that was validated using NMR spectroscopy. The PIP-binding site was located on a highly dynamic α-helix that also functions in RNA binding. Broad PIP-binding activity was observed in solution using a fluorescence polarization assay or in the context of a lipid bilayer using an on-chip, fluorescence assay. All-atom molecular dynamics simulations of the 3C protein-membrane interface revealed PIP clustering and perhaps PIP-dependent conformations. PIP clustering was mediated by interaction with residues that interact with the RNA phosphodiester backbone. We conclude that 3C binding to membranes will be determined by PIP abundance. We suggest that the duality of function observed for 3C may extend to RNA-binding proteins of other viruses.

**Highlights:**

- A viral PIP-binding site identified, validated and characterized
- PIP-binding site overlaps the known RNA-binding site
- PIP-binding site clusters PIPs and perhaps regulates conformation and function
- Duality of PIP- and RNA-binding sites may extend to other viruses

**In Brief:** The absence of conventional PIP-binding domains in viral proteins suggests unique structural solutions to this problem. Shengjuler et al. show that a viral RNA-binding site can be repurposed for PIP binding. PIP clustering can be achieved. The nature of the PIP may regulate protein conformation.

## INTRODUCTION

Positive-strand RNA viruses of eukaryotes often replicate their genomes in association with membranes (Belov and van Kuppeveld, 2012). The reason for this circumstance is not known. However, some suggest that assembly on membranes facilitates co-localization of components of viral machinery required for genome replication and virus assembly. This scenario might also conceal viral replication intermediates, for example double-stranded RNA, and progeny genomes, which are often uncapped, from interactions with cellular pattern recognition receptors and other antiviral mechanisms that would induce interferon and limit virus spread (den Boon and Ahlquist, 2010; Miller and Krijnse-Locker, 2008; Romero-Brey and Bartenschlager, 2014).

Sites of viral genome replication are created by commandeering cellular membranes. Replication compartments can be created by using organellar membranes, by hijacking vesicles of the secretory and/or endocytic pathways, or even by inducing unique membranous structures using a blend of the first two mechanisms (Belov and Sztul, 2014; Belov and van Kuppeveld, 2012; den Boon and Ahlquist, 2010; Romero-Brey and Bartenschlager, 2014; Xu and Nagy, 2014). Regardless of the origin of the membranes used for genome replication, these sites are often remodeled to produce an environment competent for maximal function. This virus-induced environment includes changes to lipid and protein composition relative to membranes found in uninfected cells.

How viral proteins are targeted specifically to sites of genome replication is a longstanding question in virology. Before 2010, conventional wisdom was that an interaction between a viral protein and a cellular protein associated with viral-induced membranes accomplished this task. Interactions between viral proteins would then recruit the full complement of the genome-replication machinery. In 2010, Altan-Bonnet and colleagues discovered that a picornavirus induced the synthesis of the phosphoinositide, phosphatidylinositol 4-phosphate (PI4P), and showed that PI4P tracked with sites of genome replication (Hsu et al., 2010). This observation inspired the hypothesis that a cellular lipid rather than a cellular protein might be the lure used to recruit viral proteins (Hsu et al., 2010).

Phosphatidylinositol-phosphate phospholipids (PIPs) function as a zip code during protein sorting to the appropriate subcellular compartment. Proteins destined for a specific cellular membrane will encode a PIP-binding determinant specific for one or a few PIPs. The most extensively characterized PIP-binding determinant is from phospholipase C (PLC), the so-called pleckstrin homology (PH) domain. This PH domain interacts with phosphatidylinositol 4,5-bisphosphate (PI(4,5)P_2_) specifically (Harlan et al., 1994). In addition to sorting, PH and related domains can be used to maintain an enzymatic activity in a latent state until the PH domain is engaged by the appropriate PIP, thus restricting enzyme activation to the appropriate subcellular compartment (Balla, 2013; Di Paolo and De Camilli, 2006; van Meer et al., 2008). All of these features are well suited to the needs of a virus to target its proteins and enzymes to the appropriate site and restrict their functions to those sites.

Do viral proteins contain PIP-binding determinants or are they targeted by association with a cellular protein containing a PIP-binding domain? Vaccinia virus, a DNA virus, is known to encode a PIP-binding protein (H7); the discovery of this protein was suggested by structural homology to the phox homology (PX) domain (Kolli et al., 2015). The retroviral matrix protein (MA) binds to PI(4,5)P_2_, which acts as a switch, promoting interaction with membrane and virus assembly (Chukkapalli et al., 2010; Saad et al., 2006). Ebola virus virion protein, VP40, also links PI(4,5)P_2_ binding to virus assembly (Johnson et al., 2016). For picornaviruses, it has been suggested that the viral polymerase binds directly to PI4P because it is retained on a PI4P-spotted hydrophobic membrane (Hsu et al., 2010). In spite of a structure for the PV polymerase and many other RNA virus-encoded proteins, structural homology to a known cellular PIP-binding protein has not been reported.

The picornaviral 3CD protein is a polyprotein cleavage product that accumulates in infected cells. This protein fuses the 3C protease domain and 3D polymerase domain. Although 3CD lacks polymerase activity, this protein contributes to many functions essential to virus multiplication (Cameron et al., 2010). 3CD interacts with membranes, induces recruitment of cellular proteins to membranes, binds to the viral genome, and somehow even stimulates virus assembly (Andino et al., 1993; Belov et al., 2007; Franco et al., 2005; Parsley et al., 1997; Pathak et al., 2007).

We used a molecular docking algorithm to identify surfaces of the PV 3C domain of 3CD with the potential to serve as sites of PIP binding. The results from computation were validated empirically by using nuclear magnetic resonance spectroscopy (NMR). The PIP-binding site overlaps the well-established RNA-binding site of PV 3C (Amero et al., 2008). We characterized the PIP-binding specificity of 3C in a soluble system as well as in the context of a lipid bilayer. Both approaches revealed broad, yet specific, binding to a combination of mono-, bis- or trisphosphorylated phosphoinositides. Binding of PV 3C to PIPs and RNA was mutually exclusive. All-atom molecular dynamics simulations of 3C interacting with PI4P- or PI(4,5)P_2_-containing membranes revealed PIP clustering and a multivalent interaction with 3C. The conformation of 3C on the membrane was modulated by the nature of the bound PIP, perhaps regulating 3C function. This model for the interaction of PV 3C with a lipid bilayer suggests that the localization of 3C domain-containing proteins is driven by PIP abundance rather than the specificity of the PIP-binding domain. The versatile use of an RNA-binding surface of a PV protein for PIP binding as well may suggest a similar duality of function in related RNA-binding proteins of other RNA viruses.

## RESULTS

### Molecular docking simulations reveal putative PIP-binding sites on PV 3C

The availability of structural data for PV 3C permitted use of a molecular-docking algorithm to screen for putative PIP-binding sites. We prepared the structure of PI4P by modifying the PI(4,5)P_2_ ligand extracted from the NMR complex structure of HIV-1 MA protein bound to PI(4,5)P_2_ (PDB ID 2H3Z) (Saad et al., 2006). The docking study of flexible PI4P with 20 active torsions to 3C protein was carried out by using the AutoDock 4.2 suite of programs. Results from one hundred 3C-PI4P docking trials predicted two clusters on 3C capable of interacting with PI4P, where 94% of the trials appeared in the major cluster and the remaining 6% appeared in the minor cluster **(Figure 1A)**. The major cluster is located near the N-terminal helix, which is positioned adjacent to a conserved loop implicated in RNA binding (Amero et al., 2008). The minor cluster is on the opposite side of the N-terminus of 3C **(Figure 1A)**.

**Figure 1.**
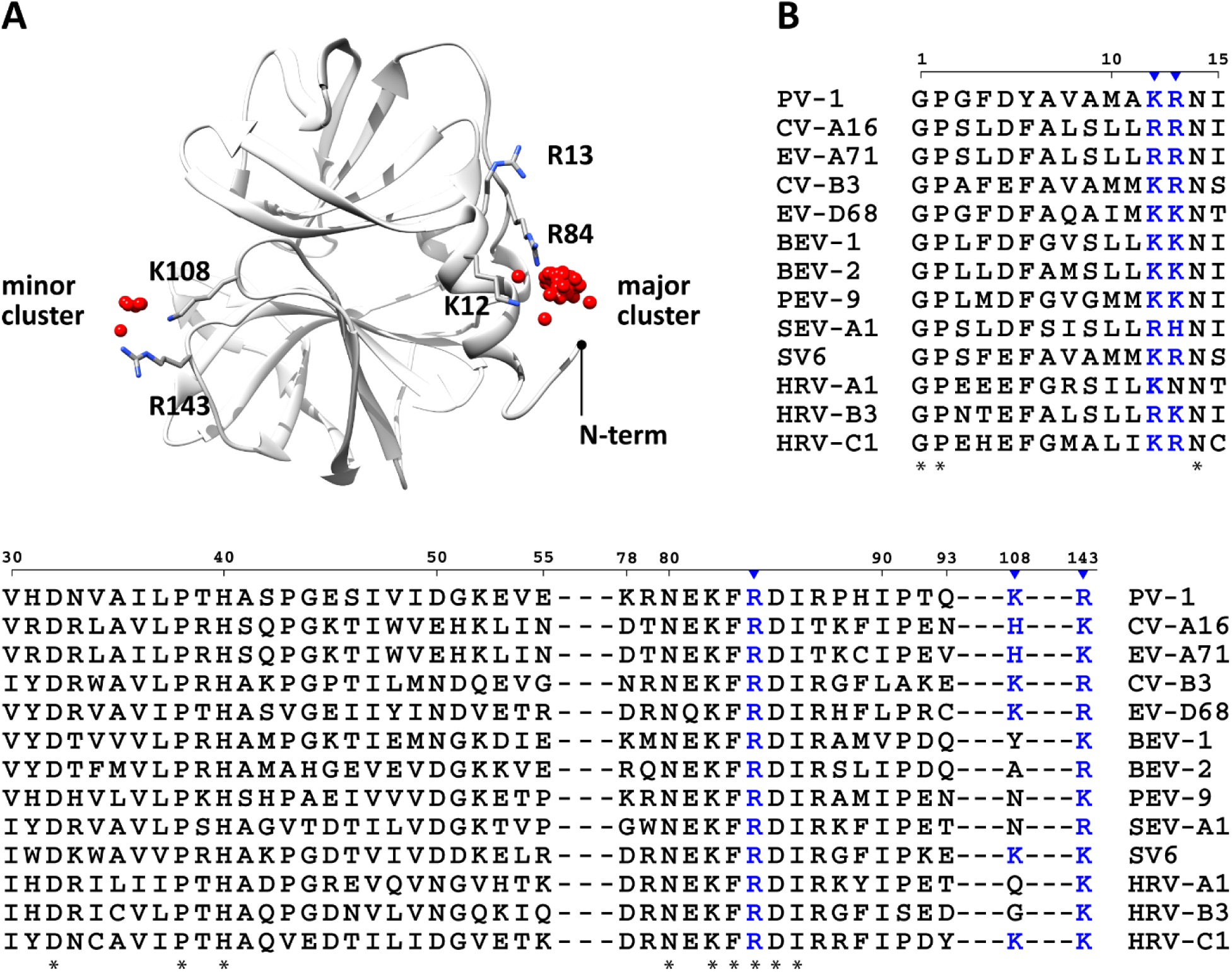
Docking analysis reveals a putative phosphatidylinositol 4-phosphate (PI4P)-binding site near the N-terminus of poliovirus 3C. **(A)** Two clusters were identified by docking dibutyl-PI4P (red spheres) on 3C surface (grey ribbon). The major cluster (94% of the trials) encompasses residues K12, R13, and R84; the minor cluster (6% of the trials) encompasses residues K108 and R143. Total of 100 docking runs were performed. The crystal structure of 3C (PDB 1L1N) was obtained from the Protein Data Bank (Mosimann et al., 1997). The structure of the PI4P ligand was prepared by modifying the structure of phosphatidylinositol 4,5-bisphosphate (PI(4,5)P_2_), extracted from the NMR complex structure of HIV-1 matrix protein bound to PI(4,5)P_2_ (Saad et al., 2006). Phosphates on the inositol head group are depicted as red spheres; basic residues in the major and minor clusters are depicted as sticks. **(B)** Multiple-sequence alignment of enteroviral 3C proteins shows that the majority of the residues predicted by docking (K12, R13, R84, and R143) and/or the basic charge are conserved; K108 is not conserved (colored in blue). Residues implicated in phosphoinositide-binding (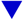); conserved residues (*); PV, poliovirus; CV, coxsackievirus; EV, enterovirus; BEV, bovine enterovirus; PEV, porcine enterovirus; SEV, simian enterovirus; SV, simian virus.

Ninety-four percent of the docking solutions in the major cluster exhibited a similar head group orientation. Red spheres indicate the position of the phosphate at position four of the inositol ring **(Figure 1A)**. In contrast, the orientation of the fatty acyl chains varied for each docking solution. Docking results indicated that K12, R13, and R84 of the major cluster interact PI4P **(Figure 1A)**. Similarly, K108 and R143 of the minor cluster are positioned to interact with PI4P **(Figure 1A)**. These residues are highly conserved in the 3C proteins of all prototype viruses in the *Enterovirus* genus of the *Picornaviridae* family **(Figure 1B)**. Even though the sequence similarity between members of the genus ranges from 40-60%, the basic nature of K12, R13, R84, and R143 were conserved, whereas that of K108 was not **(Figures 1B)**.

### Validation of PIP-binding sites by NMR

In order to test the validity of our docking observations, we titrated ^15^N-labeled 3C protein with soluble dibutyl-PI4P to observe potential NMR chemical shift perturbations (CSPs), which would indicate chemical environment changes in the presence of PI4P (**Figure 2**). Out of the three basic residues of the major cluster that were predicted to interact with the PI4P, R13 showed the largest CSP **(Figure 2A)**. There were also smaller CSPs associated with K12 and R84 (**Figure 2A**). However, we did not observe any CSPs for the minor cluster residues K108 and R143.

**Figure 2.**
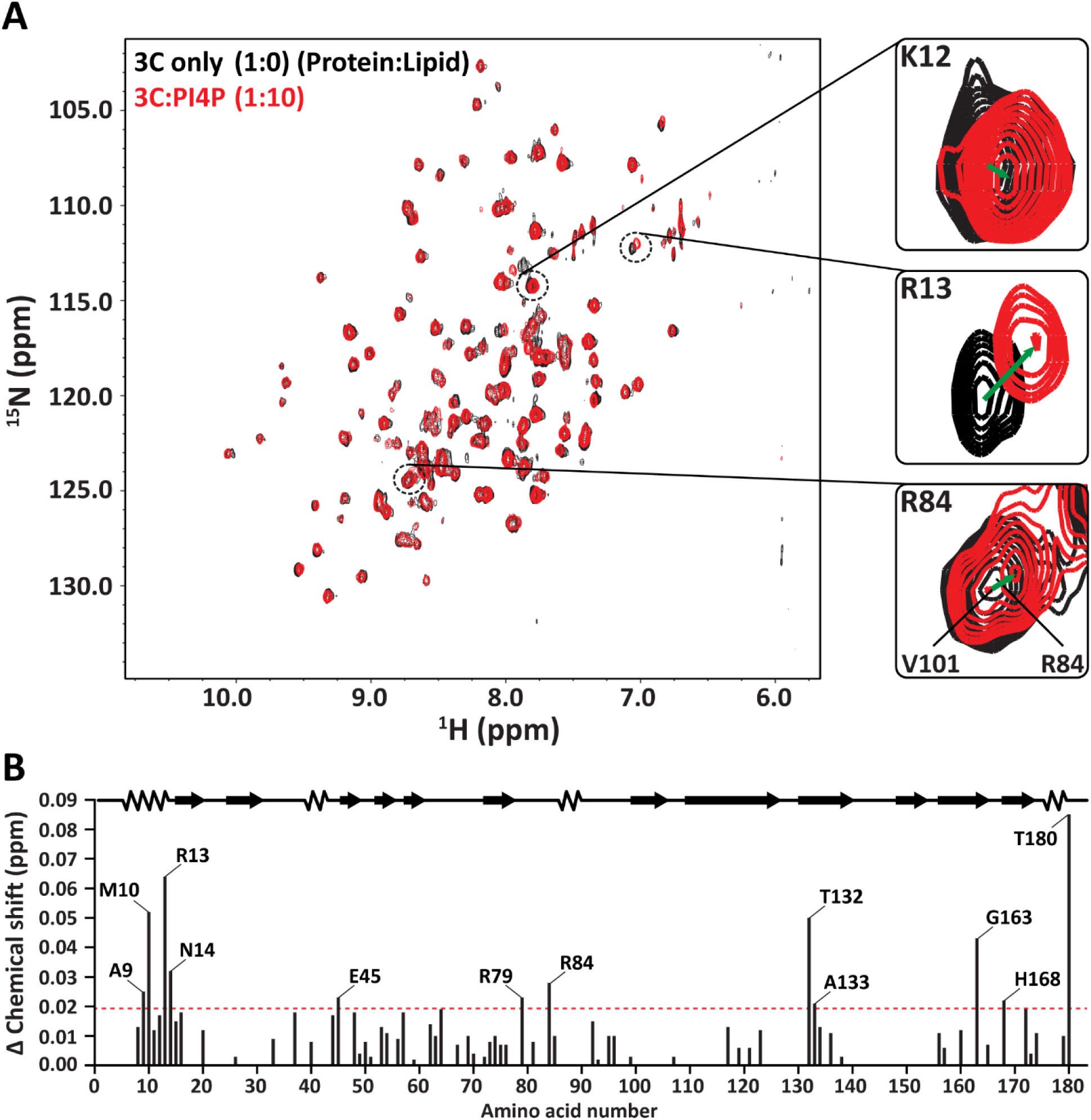
NMR spectral perturbations reveal the interaction between 3C and PI4P. **(A)** An overlay of ^1^H-^15^N heteronuclear single quantum correlation (HSQC) spectra of free 3C (black) and 3C in a complex with dibutyl-PI4P (red). (Right) Close up view of the chemical shift perturbations (CSPs) for K12, R13, and R84, which are implicated in PI4P-binding by docking. The NMR experiments were conducted at 25°C in a buffer containing 10 mM HEPES at pH 8.0, 50 mM NaCl, and 10% D_2_O. The final concentration of 3C and PI4P were 0.2 mM and 2.0 mM, respectively. **(B)** A bar graph showing CSPs induced throughout 3C by PI4P binding. The total chemical shift change (Δδ_total_) depends on the chemical shift changes in the ^1^H (Δδ_H_) and ^15^N (Δδ_N_) dimensions according to Δδ_total_ = (Δδ_H_^2^ + 0.2 Δδ_N_^2^)^0.5^. Residues with significant CSPs (above the red line) are indicated. CSPs at least one standard deviation above the average were considered to be substantial (Δδ_total_ = 0.019 ppm). These results are in agreement with the docking analyses, which show PI4P-binding towards the N-terminus of 3C.

As seen in **Figure 2B**, the addition of PI4P resulted in a number of CSPs beyond that of K12, R13, and R84. CSPs at least one standard deviation above the mean were considered to be significantly different (CSP >0.019 ppm). Many of these residues were not in the immediate vicinity of a docked PI4P molecule. For example, A9, M10, and N14 are located at the N-terminus of 3C at a distance from PI4P but all showed significant CSPs **(Figure 2B)**. These and other perturbations could be caused by other structural changes conferred by PI4P binding.

### Broad PIP-binding activity of 3C observed in solution

In order to assess PIP binding to 3C empirically and evaluate specificity of any observed binding, we used a fluorescence polarization (FP)-based PIP-binding assay (Ceccarelli et al., 2007; Kolli et al., 2015). This assay employed water-soluble dihexyl-PIPs that are BODIPY-FL-labeled at the *sn*-1 position **(Figure 3A)**. Once excited by plane-polarized light, the unbound form of the fluorescent ligand depolarizes the emitted light due to its small size and rapid rotation. In contrast, the bound form of the fluorescent ligand has a larger molecular volume, which reduces the rotation such that the emitted light remains in the same plane as that used for excitation for a longer period of time **(Figure 3A)**. This approach was validated by using the PLC-δ1 PH domain as a positive control, which showed PI(4,5)P_2_ binding specificity, as expected **(Figure S1)** (Harlan et al., 1994).

**Figure 3.**
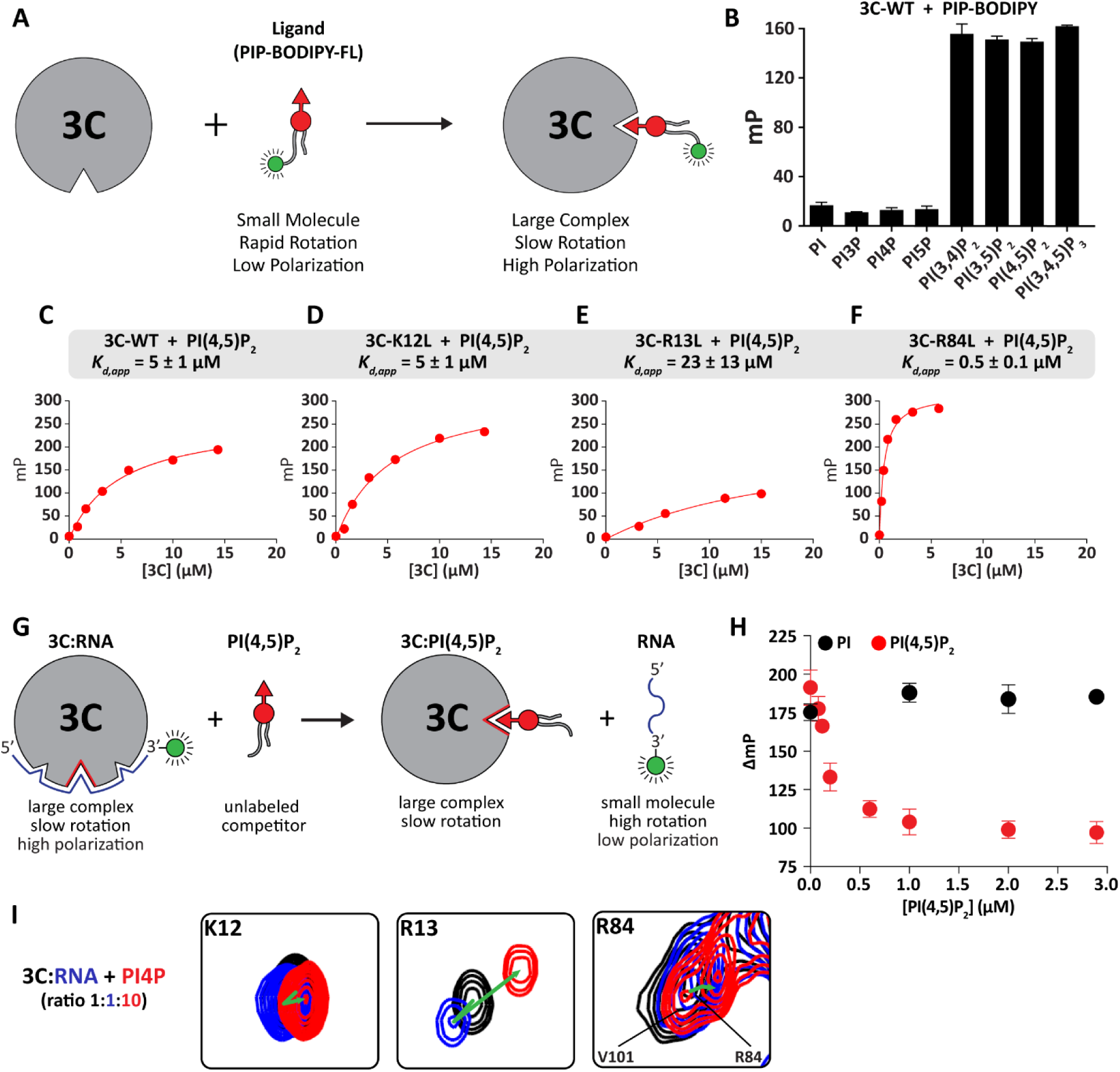
3C binds to phosphoinositides (PIPs) with broad specificity, and PIP binding competes with RNA binding. **(A)** The principles of the fluorescence polarization (FP)-based PIP-binding assay. Unbound form of the fluorescent ligand (PIP-BODIPY-FL) yields low polarization value due to its small size and rapid rotation. In contrast, the bound form yields a higher polarization value due to the overall size of the complex and its slow rotation. All FP-based PIP-binding experiments were conducted in a buffer containing 20 mM HEPES at pH 7.5 and 10 mM NaCl. 3C-PI(4,5)P_2_ samples were incubated for 30 s inside a chamber at 25 °C prior to measuring the millipolarization (mP). **(B)** PIP-binding specificity of 3C. 3C binds to both bis- and tris-phosphorylated PIP species. Binding was tested at a fixed 3C concentration (5 µM) using 0.4 nM of each PIP-probe as indicated. Error bars represent the SEM (n = 3). **(C)** WT 3C and 3C variants: **(D)** K12L, **(E)** R13L, and **(F)** R84L bind to phosphatidylinositol 4,5-bisphosphate (PI(4,5)P_2_) with varying affinities. Shown is a hyperbolic fit for each data set from which each apparent dissociation constant (*K_d,app_*) was obtained. Values are provided in **Table 1. (G)** The principles of the fluorescence polarization (FP)-based competition assay. 3C-RNA (fluorescently-labeled 11-mer) complex is pre-formed, which yields a high polarization value. As the unlabeled dibutyl-phosphatidylinositol 4,5-bisphosphate (PI(4,5)P_2_) competes out the bound RNA, the millipolarization (mP) value decreases. **(H)** FP-based competition experiments. The unlabeled PI(4,5)P_2_ efficiently competes out the bound RNA, suggesting that these interactions are mutually exclusive. The competition experiment with phosphatidylinositol (PI) competitor is used to show that binding to 3C is required for RNA displacement. Binding reactions contained 1 µM 3C-WT, 0.4 nM 3′-fluorescein (FL)-labeled RNA. Error bars represent the SEM (n = 3). **(I)** NMR-based competition experiments. 3C only (black); 3C-RNA complex (blue); 3C-PI4P complex (red). Changes in the direction of K12, R13, and R84 resonances (green arrow) in two-dimensional space suggest that RNA and PIP interactions are mutually exclusive. The NMR-based experiments were conducted at 25°C in a solution containing 10 mM HEPES at pH 8.0, 50 mM NaCl, and 10% D_2_O. The final concentration of 3C, RNA, and PI4P were 0.2 mM, 0.2 mM, and 2.0 mM, respectively.

We measured binding of phosphatidylinositol (PI) and mono-, bis-, or tris-phosphorylated PIPs to 3C **(Figure 3B)**. Binding of PI and all mono-phosphorylated PIPs was very weak. However, binding of bis- and tris-phosphorylated PIPs was substantially stronger and without any apparent order or preference **(Figure 3B)**. Given the absence of an observed binding preference of 3C, we determined the apparent dissociation constant (*K_d,app_*) for PI(4,5)P_2_ binding to 3C. The value was 5 ± 1 µM **(Figure 3C)**.

To assess the contribution of 3C residues implicated in PIP-binding to the actual binding event monitored computationally (**Figure 1A**), we changed K12, R13, or R84 to Leu. We used Leu instead of Ala to limit any collateral damage that might be caused by creating a void in the protein that could be filled by water. One expectation was that loss of any of these residues would diminish PIP binding. However, we observed that the K12L derivative behaved as WT (*K_d,app_* = 5 ± 1 µM) **(Figure 3D)**. The R13L derivative reduced binding by ~4-fold relative to WT (*K_d,app_* = 23 ± 14 µM) **(Figure 3E)**. The R84L derivative actually increased PIP-binding affinity by ~10-fold (*K_d,app_* = 0.5 ± 0.1 µM) **(Figure 3F)**.

### Binding of PIP and RNA to 3C is mutually exclusive

The residues of 3C implicated in PIP-binding (**Figures 1A** and **2B**) overlap with those reported to be required for RNA binding (Amero et al., 2008). Reported affinities of 3C for RNA are in the low micromolar range (Pathak et al., 2007), which is on par with that for PIPs. Therefore, it is likely that RNA- and PIP-binding activities of 3C will compete. To test this possibility directly, we used the FP assay to ask if the polarization of a 3C-RNA (3′-fluorescently labeled RNA) could be displaced by water-soluble dibutyl-PI(4,5)P_2_ (**Figure 3G**). We observed a concentration-dependent reduction in polarization (**Figure 3H**) that required PIP binding for displacement as dibutyl-PI exhibited no effect on RNA binding (**Figure 3H**). We also performed an NMR-based competition experiment. Because dibutyl-PI(4,5)P_2_ caused precipitation during the NMR experiment, dibutyl-PI4P was used instead. We monitored residues that were implicated in PIP binding: K12, R13, and R84. It should be noted that the CSPs induced by addition of PI4P (**Figure 2A**) or RNA (**Figure 3I**) exhibited different trajectories. The addition of PI4P to the 3C-RNA complex (**Figure 3I**) led to resonance positions similar to that of the 3C-PI4P complex (**Figure 2A**), suggesting that PI4P competed out RNA and that PI4P and RNA binding were mutually exclusive.

### Broad PIP-binding activity of 3C observed on membranes

It is possible that the context in which a PIP is presented contributes to binding affinity and/or specificity. To address this question, we employed a pH-modulation assay to monitor protein binding to a planar supported lipid bilayer (SLB) in a microfluidic device (Jung et al., 2009; Shengjuler et al., 2017). The SLB is a stable model membrane system, and the composition of the membrane can be easily tailored to probe interactions of specific lipids with proteins. The SLB used here employed three lipids, the structures of which are illustrated in **Figure 4A**. A pH-sensitive dye, *ortho*-Sulforhodamine B-POPE (*o*SRB-POPE), is embedded into the SLB and used to probe protein binding by detecting changes in the local electric field **(Figures 4B)**. Upon binding of 3C, which is positively charged at the experimental pH, the interfacial potential increases and more hydroxide ions are recruited locally. As a result, the fluorescence signal decreases **(Figures 4B and S2)**.

**Figure 4.**
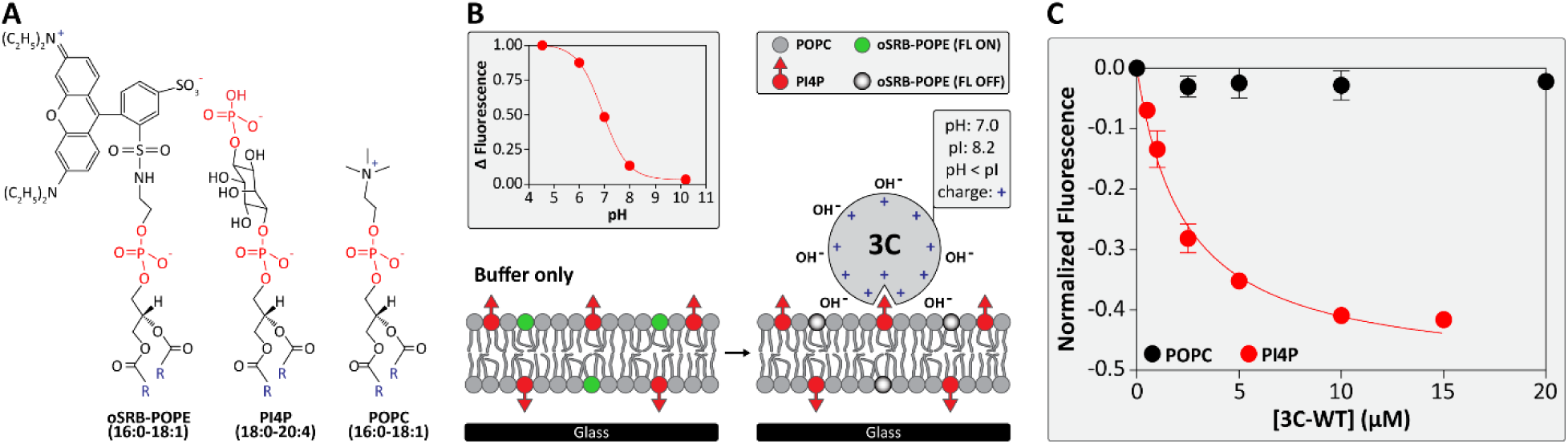
Binding of a monophosphorylated PIP to 3C in the context of a membrane. **(A)** Chemical structures of the supported lipid bilayer (SLB) components; pH-sensitive *ortho*-Sulforhodamine B-1-palmitoyl-2-oleoyl-*sn*-glycero-3-phosphoethanolamine (*o*SRB-POPE); L-α-phosphatidylinositol 4-phosphate (PI4P), and 1-palmitoyl-2-oleoyl-*sn*-glycero-3-phosphocholine (POPC). “R” represents the fatty acyl chains. **(B)** Schematic diagram illustrating the principles of the SLB-binding experiment. In the absence of 3C, the fluorescent probe is in its "on state" (left). Upon binding to the membrane, the interfacial potential is increased, causing the fluorescent probe to switch to its "off state" (right). The pH response curve of the fluorescent probe in a bilayer containing 92 mol% POPC, 7.5 mol% PI4P, and 0.5 mol% *o*SRB-POPE is shown. **(C)** 3C binding to PI4P-containing SLBs. Change in fluorescence intensity was observed as a function of 3C concentration, which was normalized to a reference channel. Shown is a hyperbolic fit of the data set. The apparent dissociation constant for 3C-PI4P interaction is 2.36 ± 0.38 µM. 3C was unable to bind to pure POPC membranes. All experiments were conducted at 20 mM HEPES at pH 7.0, 100 mM NaCl, and 5 mM magnesium acetate. Error bars represent the SEM (n = 3).

In these studies, we used 1-palmitoyl-2-oleoyl-*sn*-glycero-3-phosphatidylcholine (POPC) as the primary component of the SLB. We performed experiments in the presence of Mg^2+^ divalent cations to better mimic the cellular environment. 3C did not bind to pure POPC membranes (**Figure 4C**). The addition of PI4P to the POPC membrane revealed the capacity of 3C to bind to this mono-phosphorylated PIP **(Figure 4C)**. The value for the *K_d,app_* was 2.4 ± 0.4 µM **(Table 1)**. In addition to PI4P, 3C bound to POPC membranes containing PI3P and PI(4,5)P_2_, all with essentially the same affinity (**Table 1**). PIP binding to 3C in the context of a membrane also required the phosphates because phosphatidylinositol (PI) binding was weak (**Table 1**). Worth noting, PI binding did occur when compared to POPC alone; however, we were unable to produce a complete binding isotherm because of the extremely high concentration of protein required. The presence of PI will produce a net negative charge on the membrane, so the interaction with the basic surface of 3C makes sense. To test the possibility that negative charge alone can promote binding of 3C to the membrane, we performed an experiment in a membrane containing 1-palmitoyl-2-oleoyl-*sn*-glycero-3-phospho-L-serine (POPS). The presence of POPS will also produce a net negative charge on the membrane. The affinity of 3C for the POPS-containing membrane was on par with that observed for the PI-containing membrane (**Table 1**).

**Table 1.** Apparent dissociation constants for 3C-WT and its variants from PIP-on-a-chip experiments

| **Lipid** | **$K_{d,app} \pm SEM$ ($\mu M$)** | **Membrane comp.** | **mol%<sup>a</sup>** |
| --- | --- | --- | --- |
| **POPC** | No Binding | POPC | 99.5 |
| **PI** | $\geq 25^b$ | POPC | 92.0 |
|  |  | PI | 7.5 |
| **PI3P** | $2.4 \pm 0.4$ | POPC | 92.0 |
|  |  | PI3P | 7.5 |
| **PI4P** | $2.4 \pm 0.4$ | POPC | 92.0 |
|  |  | PI4P | 7.5 |
| **PI(4,5)P<sub>2</sub>** | $2.0 \pm 0.3$ | POPC | 92.0 |
|  |  | PI(4,5)P<sub>2</sub> | 7.5 |
| **POPS** | $\geq 47^b$ | POPC | 92.0 |
|  |  | POPS | 7.5 |
| **$K_{d,app} \pm SEM$ ($\mu M$)** |  |  |  |
| **Lipid** | **3C-K12L** | **3C-R13L** | **3C-R84L** |
| **PI4P** | $1.4 \pm 0.1$ | $4.7 \pm 0.5$ | $0.3 \pm 0.1$ |
| **PI(4,5)P<sub>2</sub>** | $1.1 \pm 0.1$ | $2.2 \pm 0.5$ | $0.3 \pm 0.1$ |

<sup>a</sup>Each membrane also contained 0.5 mol% oSRB-POPE.
<sup>b</sup>PI and POPS binding as a function of 3C concentration was in the linear range. Assuming an amplitude similar to that of PI4P binding, yields the indicated values for $K_{d,app}$ , which represent a lower limit.

### 3C-induced clustering of PIPs and PIP-induced conformations of 3C observed by MD simulations

The availability of the Anton supercomputer made it possible to perform an all-atom molecular dynamics simulation (MD) of the interaction of PV 3C with a POPC bilayer containing either PI4P or PI(4,5)P_2_ in an aqueous environment containing Na^1+^ and Cl^1-^ ions (Li and Buck, 2017b). The simulation was run for 400 ns, and the last 300 ns of the trajectory was used for analysis. The major findings of the simulation are illustrated in the panels of **Figure 5**. Binding of 3C induced clustering of at least five PIP molecules in the bilayer, independent of the nature of the PIP used (**Figures 5A and 5B**). Although the same residues of 3C were used to interact with both PIPs, the orientation of 3C differed slightly when bound to PI4P compared to that bound to PI(4,5)P_2_ (**Figures 5A and 5B**). For reference, residues of the protease active site have been indicated by black spheres. The orientation of the active site in **Figure 5A** is clearly different than observed in **Figure 5B**. This analysis expanded the number of residues interacting with PIP headgroups to include residues in regions observed by NMR, for example D32, K156, and R176 (**Figures 5A and 5B**). Interestingly, the contribution of D32 to 3C-PIP interface was mediated by two Na^1+^ ions in the case of PI4P (**Figure 5A**) and one Na^1+^ ion in the case of PI(4,5)P_2_ (**Figure 5B**). A surface representation of 3C interacting with PIPs in the context of the membrane showed that 3C is propelled above the membrane by the headgroups (**Figures 5C and 5D**). The N-terminus of 3C was an exception. Rotation of 3C to bring the N-terminus into view showed that the N-terminus has a more substantive interaction with the outer leaflet of the bilayer, albeit of a non-specific nature (**Figures 5E and 5F**). The conformation and interaction of the N-terminus with membrane was also influenced by the PIP present (**Figures 5E and 5F**).

**Figure 5.**
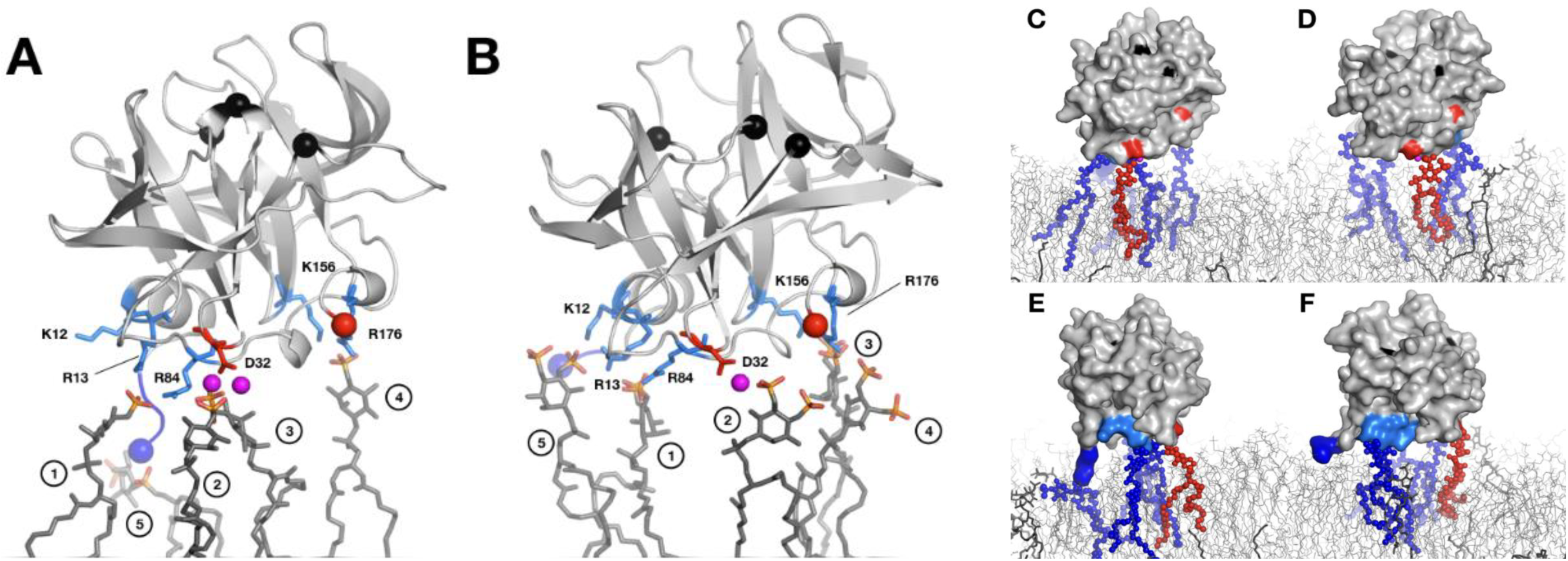
3C interacts with multiple PI4P and PI(4,5)P_2_ ligands. All-atom molecular dynamics simulations of **(A)** 3C binding to a PI4P-containing membrane or **(B)** to a PI(4,5)P_2_-containing membrane. Snapshots of selected simulations show that 3C interacts with five, clustered PI4P or PI(4,5)P_2_ molecules. PI4P and PI(4,5)P_2_ are shown as dark grey sticks with phosphates colored orange (phosphorus) and red (oxygen). 3C is shown as a light grey ribbon; residue side chains are colored as follows: light blue (positive side chains) and red (negative side chains) sticks. Also shown are: sodium ions as magenta spheres; the amino- and carboxy-termini as blue and red spheres, respectively; 3C protease active-site residues as black spheres. Specific interactions are as follows, with number in parentheses referring to headgroup in panel: (1) R13 and R84; (2) D32, mediated by sodium ions; (3) D32 for PI4P in panel A mediated by sodium, or R176 for PI(4,5)P_2_ in panel B; (4) K156 and R176; (5) α-amino group of G1 for PI4P in panel A, or K12 and R13 for PI(4,5)P_2_ in panel B. **(C)** and **(D)** Surface representations of 3C (light grey) with PI4P-containing membrane in panel C or PI(4,5)P_2_-containing membrane in panel D (blue, red and dark grey sticks); phosphatidylcholine (light grey lines). Charged interactions between 3C and the respective ligands are shown as blue (positive) or red (negative) spheres to correspondingly colored sticks. **(E)** and **(F)** same as panels **(C)** and **(D)** rotated 90 degrees counter-clockwise about the vertical access of the page to reveal the amino terminus (blue) interacting with the membrane (light grey lines). PI4P-containing membrane was composed of 244 phosphatidylcholine (PC) and 20 PI4P lipids; PI(4,5)P_2_-containing membrane was composed of 244 PC and 20 PI(4,5)P_2_ lipids. All lipids were equally distributed on each monolayer of the membrane. The simulation was conducted at 310K for 400 ns; the last 300 ns of the trajectories were used for analysis.

### SAXS reveals flexibility of N-terminus of PV 3C

For the N-terminal residues of 3C to exhibit such conformational flexibility and diverse interactions with membranes, the solution behavior would have to be more dynamic than predicted by the crystal structure (Mosimann et al., 1997). In order to evaluate the solution beavior of 3C, we performed small-angle X-ray scattering (SAXS) experiments. **Figure 6A** shows the scaled scattering profiles of 3C at concentrations ranging from 3.2 mg/mL to 13.0 mg/mL. The overlap of the scaled curves at the low ends of scattering angles was consistent with the protein remaining monomeric over this concentration range in agreement with the NMR and DLS studies (Chan et al., 2016). The scattering data at the low-angle end showed a linear correlation, satisfying the Guinier approximation (qRg < 1.3), from which we determined a radius of gyration (Rg) of 17.26 Å **(Figure 6A)**. The Rg derived from the Guinier approximation was in good agreement with that estimated from the pair-distance distribution function P(r) calculated by GNOM (real-space Rg = 17.72 Å, reciprocal-space Rg = 17.76 ± 0.13 Å) **(Figure 6B)**. From the calculated P(r), the maximum particle dimension of the protein was estimated to be 65 Å **(Figure 6B)**. The asymmetric shape of the distribution function suggested the presence of a tail in the solution structure of 3C. The values of Rg and Dmax from SAXS data also suggested that the protein exists as a monomer in solution. Furthermore, the estimated molecular mass of 22.2 ± 0.9 kDa (derived from Porod volume) or 20.5 ± 1.1 kDa (derived from SAXS MoW calculation) was in very good agreement with the calculated molecular mass of a monomer (19.6 kDa).

**Figure 6.**
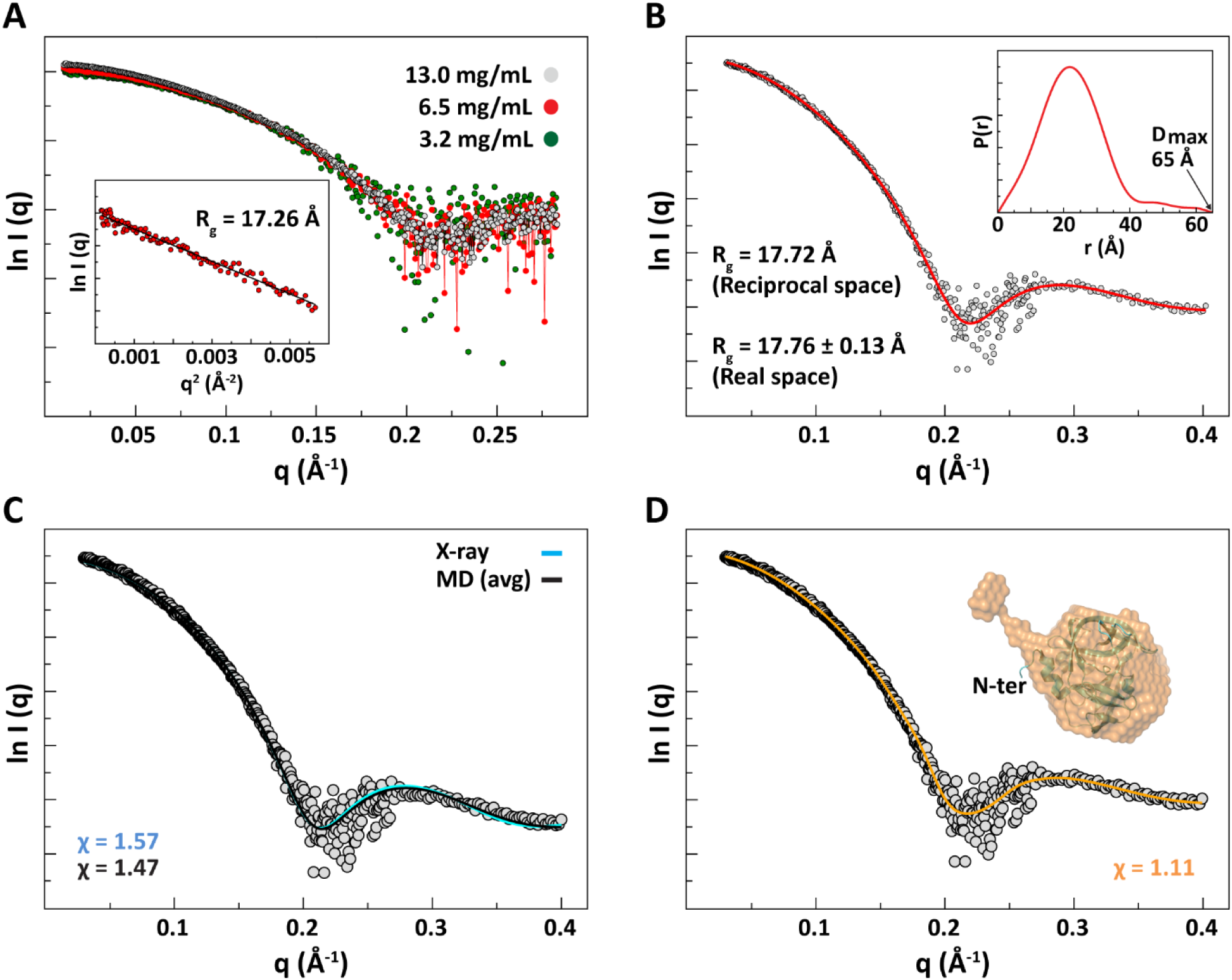
The N-terminus of 3C is dynamic as revealed by SAXS. **(A)** Scattering profiles of 3C obtained at three different protein concentrations (3.2 mg/mL, green; 6.5 mg/mL, red; 13.0 mg/mL, grey). Scaling factors were applied for the low-concentration data. The inset shows the linear fitting of the Guinier plot from which radius of gyration (R_g_) was determined. **(B)** The transformed scattering profile calculated by GNOM from the pair-distance-distribution function, P_(r)_, that is shown in the inset. The maximum particle dimension and the R_g_ obtained from P_(r)_ are indicated. The estimated R_g_ from GNOM is in good agreement with that obtained from Guinier fitting. **(C)** The calculated scattering profiles of the crystal structure of poliovirus 3C monomer (cyan) (Mosimann et al., 1997) and the average MD structure (black) (Moustafa et al., 2015) fitted to the experimental SAXS data (grey). Both structures fit well; however, the average MD structure showed slightly better fitting compared to the crystal structure as indicated by its lower χ-value. **(D)** The reconstructed SAXS filtered model calculated by DAMMIN is shown as an orange transparent surface with the crystal structure superimposed. The SAXS model clearly shows an extended N-terminus compared to the crystal structure. The calculated scattering profile of the SAXS model (orange) fitted to the experimental data (grey) is also shown.

We compared the calculated scattering profiles of the crystal structure and the average MD structure to the experimental data. As shown in **Figure 6C**, both structures agreed well with the experimental data. However, the calculated curve from the average MD structure showed a slightly better fitting to the SAXS data relative to the crystal structure (χ = 1.57 for crystal structure vs. χ = 1.47 for MD structure). Next, we built the *ab initio* low-resolution SAXS model using DAMMIN and DAMMIF programs; 10 independent models from each program were generated and averaged. The models were highly similar; the average values of the normalized spatial discrepancy (NSD), which represents the similarity among the models, were 0.563 ± 0.01 and 0.698 ± 0.064 for DAMMIN and DAMMIF models, respectively. Superimposing the average MD structure onto the SAXS model showed a very good agreement between these two models with an NSD of 0.66. The crystal structure showed less agreement with the SAXS model with an NSD of 0.92. The reconstructed low-resolution SAXS model clearly revealed the presence of an extended and highly dynamic N-terminal helix of 3C, the volume of which exceeded the volume of a helix that is 14 residues in length **(Figure 6D)**. Worth noting, the amphipathic nature of this helix (**Figure S3**) may also contribute to the alternative conformations observed on membranes (**Figures 5E and 5F**).

## DISCUSSION

The membranes forming the genome-replication organelle of enteroviruses are enriched in PI4P. It is known that the RNA-dependent RNA polymerase domain, 3D, of enteroviruses binds to PI4P, although it is not clear where on the protein this binding occurs (Hsu et al., 2010). Because the 3CD protein accumulates to a much higher level in PV-infected cells than 3D and 3CD has been implicated in formation and/or function of the genome-replication organelle (Oh et al., 2009), we used the available structural information for these proteins (Marcotte et al., 2007; Mosimann et al., 1997; Thompson and Peersen, 2004) to screen for PI4P-binding sites by using a molecular-docking algorithm (Goodsell et al., 1996; Morris et al., 1998). The observation of a putative PI4P-binding site on 3C was unexpected but exciting as it was now very possible that trafficking of PV proteins to the site of genome replication might be facilitated by use of new structural classes of PI4P-binding domains (**Figure 1**). Our focus on 3C was motivated by the arsenal of structural and computational resources available to study PV 3C that would facilitate identification and characterization of the PI4P-binding site (Amero et al., 2008; Marcotte et al., 2007; Mosimann et al., 1997; Moustafa et al., 2015), including a near-complete backbone NMR resonance assignment of the 2D (Amero et al., 2008).

Using a solution- and bilayer-based assay for PIP binding, we observed PIP binding by 3C (**Figures 3** and **4**). Titration of PI4P into a solution containing 3C caused CSPs that were consistent with the major PI4P-binding site observed computationally (**Figure 2A**). However, the CSPs also included other residues (**Figure 2B**), in particular residues known to contribute to the RNA-binding activity of 3C (Amero et al., 2008). PIP-binding was indeed competitive with RNA binding (**Figures 3H and 3I**). For a small molecule like PI4P to cause so many CSPs or to compete with a substantially larger RNA, either multiple molecules of PI4P bind to the protein or a single molecule of PI4P induces a large change in the conformation and/or dynamics of the protein. Analysis of NMR spectra were inconsistent with large-scale changes in conformation or dynamics as most of the resonance positions and intensities were not affected by PI4P binding (**Figure 2**). MD simulations were consistent with multiple PIPs binding to PV 3C along the shallow cleft used for RNA binding (**Figure 5**). While the negative charge of the phospholipid headgroup contributed to PIP binding, the display of the charge on the inositol ring contributed to high-affinity binding (**Table 1**). In this regard, neither the location of the phosphate on the inositol ring nor the number of phosphates on the inositol ring mattered for 3C binding (**Table 1**), although the nature of the PIP influenced the conformation of 3C bound to membrane observed computationally (**Figure 5**). MD simulations indicated that one contributor to the PIP-dependent conformation is the flexibility of the N-terminus (compare **Figures 5E and 5F**). Such conformational flexibility of the 3C N-terminus was also observed in solution using SAXS (**Figure 6**).

The PIP-binding site structure, corresponding specificity and extreme capacity to cluster PIPs differs substantially from known cellular PIP-binding domains. Most cellular PIP-binding domains contain a solvent-accessible, basic cavity to which a PIP binds (Lemmon, 2008). In certain cases, spatial restriction within the cavity enables stereospecificity to the interaction with a PIP headgroup (Lemmon, 2008). Even though the putative PIP-binding sites on 3C are enriched in basic residues, the surface is shallow by comparison to cellular PIP-binding domains. While cellular PIP-binding proteins are known to bind more than one PIP or a single PIP in combination with another anionic phospholipid, the maximum number is usually two PIPs (Moravcevic et al., 2012; Yamamoto et al., 2016). The expanded specificity does not represent mere sloppiness but permits regulation by the simultaneous presence of multiple PIPs, which has been referred to as coincidence detection (Moravcevic et al., 2012). The broad specificity of the 3C PIP-binding site is unprecedented in mammalian systems but broad specificity may be more common in the yeast, *Saccharomyces cerevisiae* (Yu et al., 2004). This circumstance may not cause a problem for the virus because PI4P synthesis is induced immediately upon infection, and the levels of other PIPs in the cell exist at a much lower level (van Meer et al., 2008). Much of the apparent affinity of 3C for the PIP-containing membrane derives from the multivalent nature of the interaction (**Figure 5**). Clustering of PIPs has been suggested by MD of cellular PIP-binding proteins bound to membranes; however, binding of two PIPs is most common (Yamamoto et al., 2016).

Binding of five PIPs to the same protein seems extraordinary (**Figure 5**). Whether or not the virus requires substantial PIP clustering and whether or not such substantial PIP clustering causes a biological response are questions that need to be addressed in the future. One class of PIP-binding proteins uses an unstructured basic domain to bind to a leaflet of the bilayer and sequester PIPs (McLaughlin et al., 2002). Amongst the best-characterized member of this class of proteins is MARCKS, myristoylated alanine-rich C kinase substrate (McLaughlin et al., 2002). PIP sequestration causes an increase in PIP concentration, which may serve to increase signaling intensity when acted upon by effectors such as phospholipases (McLaughlin et al., 2002). The process of sequestration can be tuned down or turned off by the presence of Ca^2+^ (McLaughlin et al., 2002). Similarly, 3C bound to a membrane should sequester PIPs. Indeed, 3C binding to membranes at any significant level could act as a phosphoinositide sink, given the presence of contact sites to connect all of the intracellular membranes (Helle et al., 2013; Raiborg et al., 2016). The ability of a virus to sequester phosphoinositides could have a profound, debilitating effect on the cell.

PIP-binding by 3C is also unique when compared to well-characterized PIP-binding proteins of RNA viruses, of which there are only two: HIV MA protein and Ebola VP40 (matrix) protein (Chukkapalli et al., 2010; Johnson et al., 2016; Saad et al., 2006). The role of viral matrix proteins is to bridge the genome or some genome-containing ribonucleoprotein complex to the membrane used to envelope the virus as it buds from the cell. Both of these viral proteins appear to bind specifically to PI(4,5)P_2_. PIP-binding signals to the protein that it has reached the destination for assembly. In the case of MA, PIP binding triggers the exposure and insertion of its myristoylated N-terminus into the plasma membrane (Saad et al., 2006). In the case of VP40, PIP binding promotes oligomerization (Johnson et al., 2016). Both HIV MA and Ebola VP40 assemble into oligomers, which cluster PIPs, but not to the same extreme as predicted for 3C (Johnson et al., 2016; Saad et al., 2006). Both of these matrix proteins bind RNA. In the case of Ebola VP40, there is no evidence to suggest that RNA competes with PIP binding (Gomis-Ruth et al., 2003). However, competition may exist for HIV MA (Chukkapalli et al., 2010).

Non-structural protein 5A (NS5A) from hepatitis C virus binds PI(4,5)P_2_ and to lesser extent PI(3,4)P_2_ through its N-terminal amphipathic α-helix (Cho et al., 2015). The motif used has been referred to as the basic amino acid PIP2 pincer (BAAPP) domain (Cho et al., 2015). This motif is nothing more than two basic amino acids flanking a series of hydrophobic residues displayed on a helix, presumably that penetrates into the bilayer (Cho et al., 2015). The BAAPP motif can be found in amphipathic α-helices of cellular and viral proteins; however, the function is not known. Indeed, such a motif might exist in the N-terminus of 3C (**Figure S3)**. In the case of 3C, we propose that this helix just augments membrane binding without any specificity (**Figures 5E and 5F**). How the BAAPP domain confers specificity is not known. It is worth noting that NS5A is an RNA-binding protein. The organization described for 3C, amphipathic α-helix followed by an RNA-binding domain, therefore applies to NS5A. It is compelling to speculate that this RNA-binding domain might also interact with PIPs.

Other virus families encode 3C-like proteases (3CL), specifically the *Caliciviridae* and *Coronaviridae*. One of the most prominent members of the Calicivirus family is Norwalk Virus and other human noroviruses (NoV), the so-called cruise-ship diarrhea viruses. Prominent members of the Coronavirus family include Severe Acute Respiratory Syndrome Virus (SARS CoV) and Middle Eastern Respiratory Syndrome Virus (MERS CoV). PV 3C, NoV 3CL (also known as NS6) and SARS/MERS CoV 3CL (also known as nsp5) share a chymotrypsin-like fold (Anand et al., 2002; Hilgenfeld, 2014; Mosimann et al., 1997; Zeitler et al., 2006). The protease active sites are quite similar. Structurally, NoV 3CL is extremely close to PV 3C, with clear conservation of residues corresponding to: K12, R13, D32, K156 and R176 (Zeitler et al., 2006). Residues of SARS/MERS CoV 3CL are not so highly conserved, but the surface of this protein corresponding to the PIP-binding surface of PV 3C is clearly basic (Anand et al., 2002). There is one report of RNA binding by NoV 3CL (Viswanathan et al., 2013) but none for the CoV proteins. The possibility that members of other virus families of medical importance encode a PIP-binding protein like that of PV 3C suggests that this activity may represent a target with broad-spectrum therapeutic potential. Empirical studies of the NoV and CoV 3CL proteins will be required to assess this possibility.

Our computational studies suggested that 3C interacted with a single PIP mediated by K12 and R84. (**Figure 1A**). Although R13 was nearby, a role for this residue in PIP binding was not suggested (**Figure 1A**). Removal of the positive charge from K12 had no impact on PIP binding, removal from R84 increased PIP-binding affinity, but removal from R13 weakened PIP-binding affinity **(Figures 3C-3F**). A role for R13 and R84 in PIP binding was supported by NMR experiments, but the evidence for K12 was weak (**Figure 2B**). So, while the molecular-docking simulation pointed us in the right direction, the molecular details of the putative interaction were not correct. The inaccuracy may have been caused by our attempt to dock a single PI4P instead of multiple PIPs. The 3C-membrane-binding simulation (**Figures 5A and 5B**) produced results that were consistent with both the NMR and mutagenesis experiments. K12 has either no or a minor role in binding PI4P (**Figure 5A**) or PI(4,5)P_2_ (**Figure 5B**). R13 interacts directly with one (**Figure 5A**) or two (**Figure 5B**) PIP phosphates. R84 is near the Na^1+^ ions at the interface; removal of the charge in this instance could be stabilizing.

The broad repertoire of PIPs bound by 3C may occur *specifically* and regulate the conformation of 3C. Although the same collection of 3C residues appear to be involved in interactions with both PI4P and PI(4,5)P_2_, the architecture of the interface is different (**Figure 5**). It is possible that these apparent states are interconvertible and just reflect the short duration of the simulated trajectory. However, it is possible that different PIPs stabilize different states. For the K-Ras4A protein, it is known that POPS stabilizes a different conformation of this protein than PI(4,5)P_2_ (Li and Buck, 2017a). During infection, 3C functions primarily in the context of a larger precursor, 3CD. 3CD is known to have many functions before, during and after genome replication (Cameron et al., 2010; Oh et al., 2017). Having functions dictated by conformational states of the protein stabilized by phospholipid composition of the membrane to which 3CD is bound would be an elegant explanation of the regulation of the many activities of this protein. Deliberate interrogation of this hypothesis is warranted.

In conclusion, in an effort to begin to identify proteins and mechanisms that PV uses to recognize PIPs, we have discovered that nature has been creative and adapted a viral RNA-binding surface to perform this additional task. The outcome is a new structural class of PIP-binding proteins with the capacity for multivalent binding to any mono-, bis- or tris-phosphorylated PIP, but with a unique conformational state determined by the nature of the bound PIP. It is likely that a similar mechanism exists in noroviruses and coronaviruses, suggesting a role for PIPs in the biology of these viruses even though empirical support of this possibility is needed.

## AUTHOR CONTRIBUTIONS

Conceptualization: CEC; Methodology: all authors; Validation: DS, YMC, SS, IMM, ZL, DWG; Formal Analysis: DS, YMC, SS, IMM, ZL, DWG; Investigation: DS, YMC, SS, IMM, ZL, DWG; Writing – Original Draft: DS and CEC; Writing – Review & Editing: all authors; Visualization: DS, YMC, IMM, DWG; Supervision: MB, PSC, DDB, CEC; Project Administration: CEC; Funding Acquisition: MB, PSC, DDB, CEC.

## ACKNOWLEDGMENTS

We thank Calvin Yeager and Jamie J. Arnold for contributing to the assembly and submission of the manuscript. Funding for this study was obtained from the following sources: CEC, AI053531 from NIAID, NIH; DDB, AI091985 and AI104878 from NIAID, NIH; PSC, N00014-14-1-0792 from Office of Naval Research; and MB, GM112491 from NIGMS, NIH.

## STAR METHODS

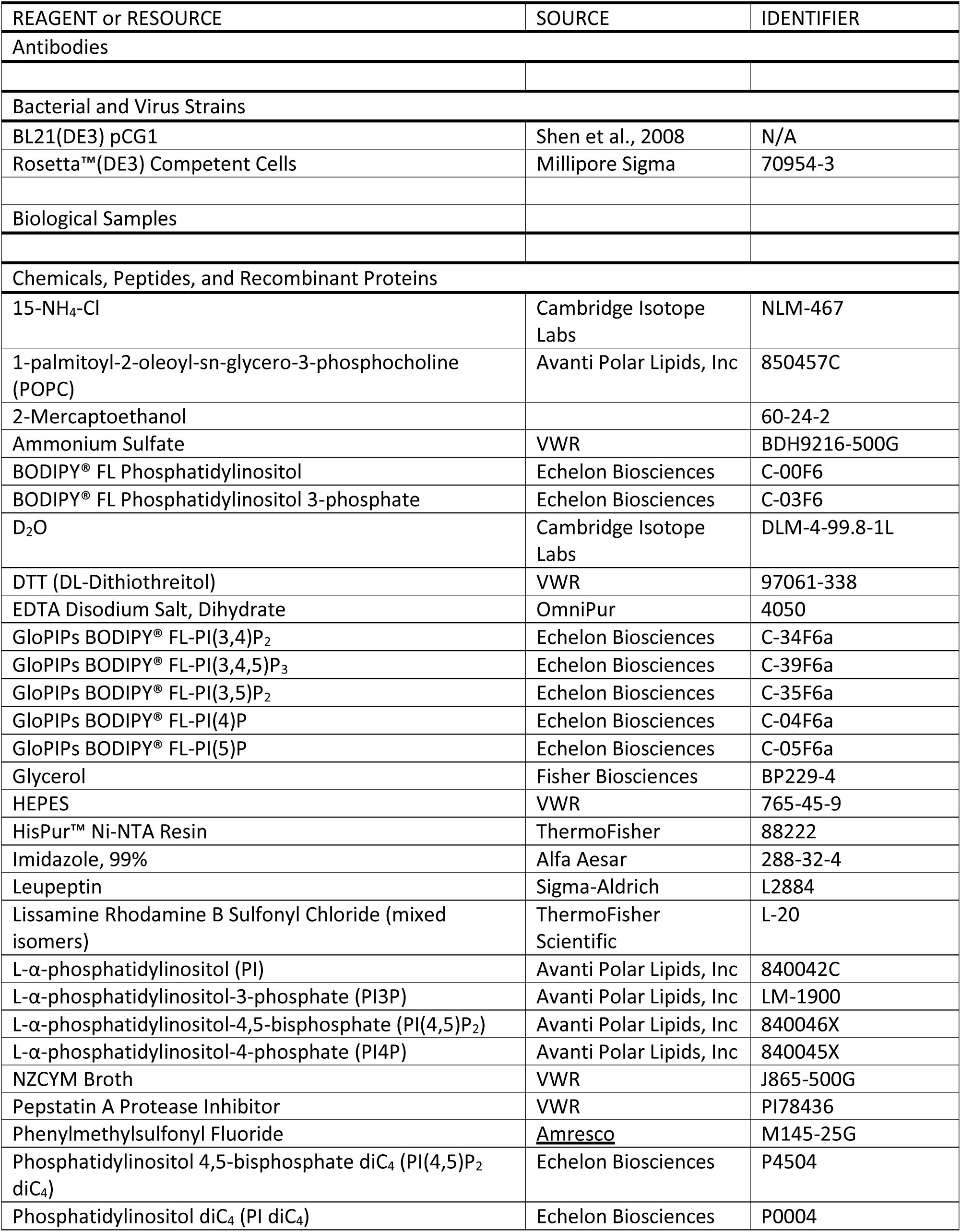

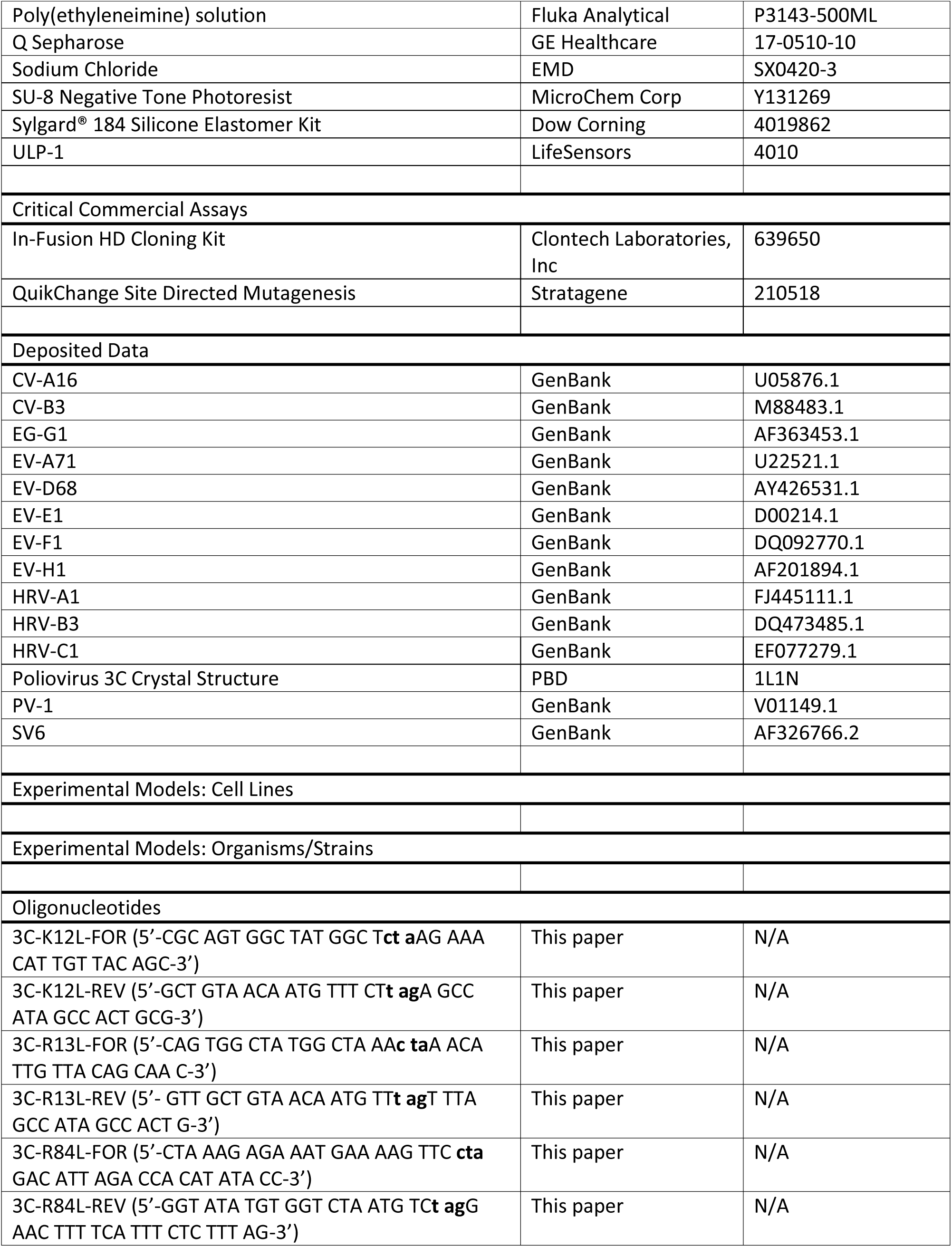

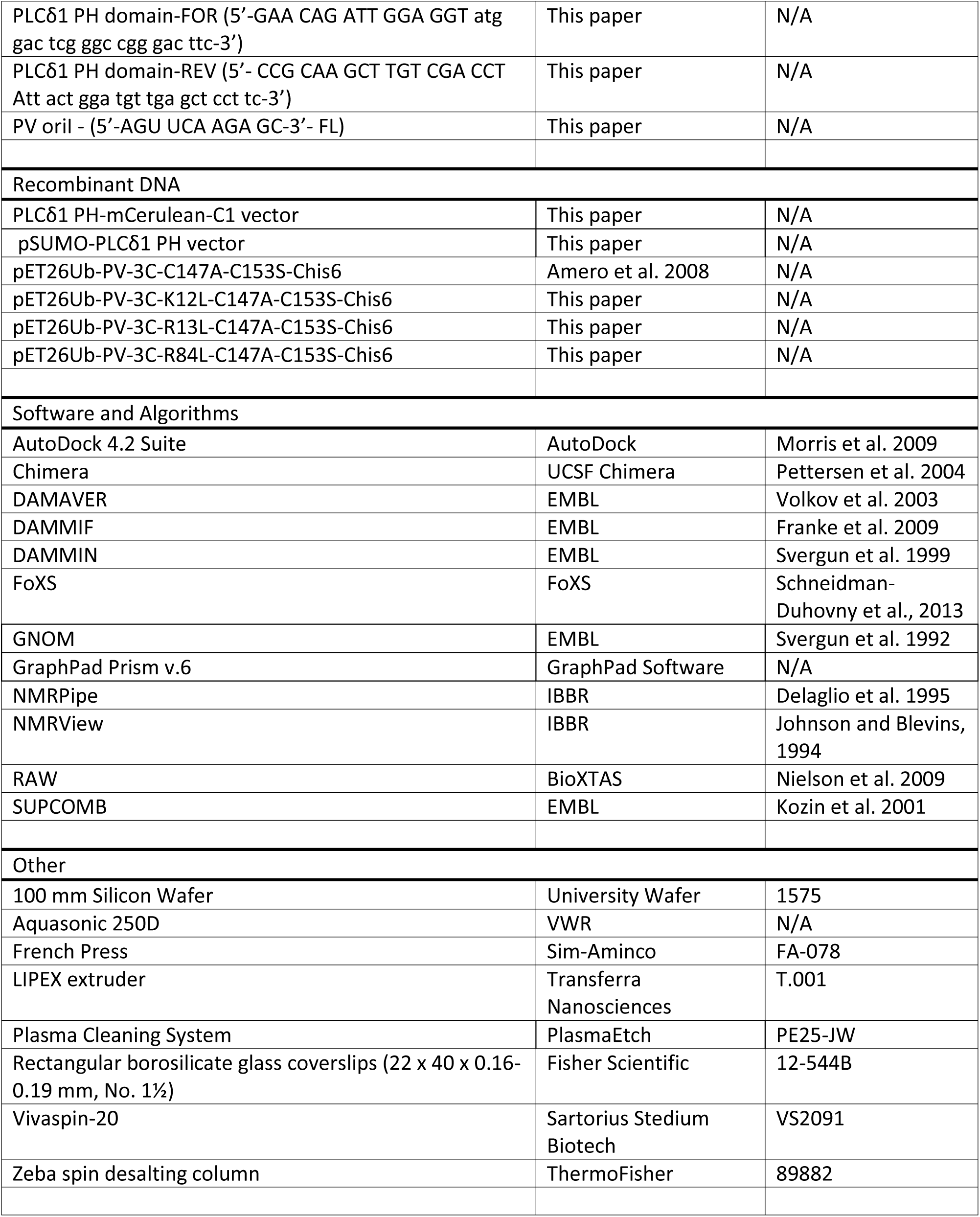

### Contact for Reagent and Resource Sharing

Further information and requests for resources and reagents should be directed to and will be fulfilled by the Lead Contact, Craig E. Cameron.

### Experimental Model and Subject Details

#### Cell Lines

Rosetta™(DE3) Competent Cells (Millipore Sigma) were used for PLC-δ1 protein purification and grown in NZCYM media, pH 7.6, by IPTG induction.

Poliovirus proteins were expressed and purified from *E. coli* BL21(DE3)pCG1 (Shen et al., 2008) in NZCYM media, pH 7.6, by IPTG induction.

### Method Details

#### Materials

BODIPY-FL-labeled and unlabeled water soluble phosphoinositides were from Echelon Biosciences, Inc. The lipids for supported lipid bilayer experiments, 1-palmitoyl-2-oleoyl-sn-glycero-3-phosphocholine (POPC) (850457C), L-α-phosphatidylinositol (PI) (840042C), L-α-phosphatidylinositol-3-phosphate (PI3P) (LM-1900), L-α-phosphatidylinositol-4-phosphate (PI4P) (840045X), and L-α-phosphatidylinositol-4,5-bisphosphate (PI(4,5)P_2_) (840046X) were from Avanti Polar Lipids. The pH sensitive fluorescent probe *ortho*-Sulforhodamine B-POPE (oSRB-POPE) was synthesized following the previously published procedure (Huang et al., 2013). ^15^N-ammonium chloride and deuterium oxide were purchased from Cambridge Isotope Laboratories. Rectangular borosilicate glass coverslips (22 x 40 x 0.16-0.19 mm, No. 1½) for supported lipid bilayer binding experiments were from Fisher Scientific (12-544B) and polydimethylsiloxane (PDMS) (Dow Corning Sylgard Silicone Elastomer-184) was from Ellsworth Adhesives (4019862). Prime grade, single side polished, silicon wafers with 100 mm diameter and 525 µm thickness were from University Wafer (ID. 1575). All other reagents were available from VWR, Fisher Scientific, or Sigma-Aldrich.

#### Construction of expression plasmids

The 3C construct used in this study, pET26Ub-PV-3C-C147A-C153S-CHis6, was previously described (Amero et al., 2008). The QuikChange site-directed mutagenesis kit (Stratagene) was used to introduce mutations into the 3C coding sequence. The oligonucleotides employed were as follows: 3C-K12L-FOR (5′-CGC AGT GGC TAT GGC T**ct a**AG AAA CAT TGT TAC AGC-3′), 3C-K12L-REV (5′-GCT GTA ACA ATG TTT CT**t ag**A GCC ATA GCC ACT GCG-3′), 3C-R13L-FOR (5′-CAG TGG CTA TGG CTA AA**c ta**A ACA TTG TTA CAG CAA C-3′), 3C-R13L-REV (5′-GTT GCT GTA ACA ATG TT**t ag**T TTA GCC ATA GCC ACT G-3′), 3C-R84L-FOR (5′-CTA AAG AGA AAT GAA AAG TTC **cta** GAC ATT AGA CCA CAT ATA CC-3′), and 3C-R84L-REV (5′-GGT ATA TGT GGT CTA ATG TC**t ag**G AAC TTT TCA TTT CTC TTT AG-3′). Bold letters indicate the codon where mutations were introduced. The sequence of each clone was confirmed by Sanger sequencing at the Penn State Genomics Core Facility in University Park, PA.

PLCδ1 PH-mCerulean-C1 vector was a kind gift of Dr. Lorraine Santy here at The Pennsylvania State University in University Park, PA. The PLCδ1 PH domain was PCR amplified by using the forward (5′-GAA CAG ATT GGA GGT atg gac tcg ggc cgg gac ttc-3′) and reverse (5′-CCG CAA GCT TGT CGA CCT Att act gga tgt tga gct cct tc-3′) primers. This creates overhangs (capital letters) on either end and introduces a Bsa I cut site at the 5′-end and Sal I cut site at the 3′-end, which was used for cloning the gene into the Bsa I and Sal I-linearized pSUMO vector via homologous recombination. In-Fusion HD cloning kit (Clontech Laboratories, Inc.) was used to generate the pSUMO-PLCδ1 PH expression plasmid.

#### Molecular Docking

The docking study of PI4P to 3C and 3D proteins was carried out using AutoDock 4.2 suite of programs (Goodsell et al., 1996; Morris et al., 1998). The crystal structure of the 3C protein (PDB 1L1N) (Mosimann et al., 1997) was obtained from the Protein Data Bank. Only chain A of the two identical monomers in the crystal structure was chosen for the docking. The protein was prepared for the docking runs as follows: the structural water molecules and any non-protein atoms were deleted from the crystal structure; explicit hydrogen atoms were added to the protein and the structure was subjected to quick minimizations (500 steps) using AMBER force field ff99SB in Chimera (Hornak et al., 2006; Pettersen et al., 2004). Next, the AutoDockTools (ADT) was used to complete the preparation of the target protein by merging the non-polar hydrogen atoms, adding Kollman charges, and creating the PDBQT files for the docking runs. The structure of the PI4P ligand was prepared by modifying the structure of PI(4,5)P_2_, extracted from the NMR complex structure of HIV-1 matrix protein bound to PI(4,5)P_2_ (PDB 2H3Z), in Chimera (Saad et al., 2006). The ligand was subjected to quick energy minimizations (100 steps) in Chimera after adding hydrogen atoms and Gasteiger-Marsili atomic partial charges; ADT was then used to prepare the ligand PDBQT file, the flexible ligand has 20 active torsions.

For the target protein, the affinity grid field was generated using the auxiliary program AutoGrid. The grid size was set to 49 x 49 x 49 points for the 3C protein with a grid spacing of 1 Å. The grid size extended at least 15 Å beyond the size of the protein in all dimensions. The docking was performed using Lamarckian Genetic Algorithm (LGA). For the target protein-ligand pair, 100 docking runs were performed with a population of 150 individuals, maximum number of 25,000,000 energy evaluations, and maximum number of 27,000 generations. The default parameters of mutation rate (0.02), crossover rate (0.80), and GA’s selection window (10 generations) were used. All other parameters were kept at their default values: translation step (2.0 Å), quaternion step (50.0), and torsion step (50.0). The 100 docking solutions resulted from the run were grouped into clusters such that the ligand root-mean-square deviation (rmsd) within each cluster were below 2.0 Å. The clusters were also ranked by the value of the lowest energy solution within each cluster.

#### Small angle X-ray scattering (SAXS) analysis

A non-His-tagged version of 3C protein was expressed and purified. Protein samples were prepared at concentrations of 3.2, 6.5, and 13.0 mg/mL in 40 mM Tris-HCl buffer at pH 7.4 containing 200 mM NaCl, 10% glycerol, 1 mM EDTA and 2 mM DTT. Synchrotron SAXS data were collected on the F2-line station at MacCHESS at 293 K using dual Pilatus 100K-S detector and a wavelength of 1.224 Å. The data were collected using exposure times of 5 minutes in ten 30-second frames and covered a momentum transfer range (q-range) of 0.01 < q < 0.8 Å^-1^. The program RAW (Nielsen et al., 2009) was used for data reduction and background subtraction. The radius of gyration (Rg) and forward scattering I(0) were calculated using Guinier approximation. The GNOM program (Svergun, 1992) was used to calculate the pair-distance distribution function P(r) from which the maximum particle dimension (Dmax) and Rg were estimated. The ab initio low-resolution models were reconstructed using DAMMIN (Svergun, 1999) and DAMMIF (Franke and Svergun, 2009) programs using data in the range 0.031 < q < 0.40 Å^-1^; ten independent models generated from each program were averaged using DAMAVER (Volkov and Svergun, 2003). FOXS (Schneidman-Duhovny et al., 2013) was used to calculate the theoretical scattering curves of the generated models and to carry out the fitting to the experimental data. SUPCOMB (Kozin and Svergun, 2001) was used to superimpose the 3C crystal structure (PDB 1L1N) to the SAXS model.

#### Sequence alignments

Polyprotein sequences of *Enterovirus* prototype strains were retrieved from GenBank; CV-A16 (U05876.1), EV-A71 (U22521.1), CV-B3 (M88483.1), PV-1 (V01149.1), EV-D68 (AY426531.1), EV-E1 (D00214.1), EV-F1 (DQ092770.1), EG-G1 (AF363453.1), EV-H1 (AF201894.1), SV6 (AF326766.2), HRV-A1 (FJ445111.1), HRVB3 (DQ473485.1), and HRV-C1 (EF077279.1). 3C sequences of each strain were aligned by Clustal Omega (www.ebi.ac.uk/Tools/msa/clustalo/) with default alignment parameters. The pre-aligned sequences were further processed by ESPript 3.0 (espript.ibcp.fr) program to generate PostScript files.

#### Expression and purification of PV 3C

3C protein was expressed in BL21(DE3)pCG1 competent cells as previously described (Shen et al., 2008). Cells were harvested by centrifugation at 5,400 x g at 4°C for 10 min and washed with a buffer containing 10 mM Tris and 1 mM EDTA at pH 8.0 and then re-centrifuged. The cell pellet was resuspended in Buffer A [20 mM HEPES, 10% glycerol, 5 mM imidazole, 5 mM β-mercaptoethanol (BME), 1 mM EDTA, 500 mM NaCl, 1.4 μg/mL pepstatin A, 1.0 μg/mL leupeptin, at pH 7.5] at a 5 mL Buffer A per 1-gram cell pellet ratio. The cell suspension was homogenized by Dounce homogenizer and lysed by passing through a French press twice at a pressure of 1,000 psi. Phenylmethylsulfonyl fluoride (PMSF) was added to the cell lysate at a final concentration of 1 mM. The suspension was clarified by centrifugation at 75,000 x g at 4°C for 30 min. Polyethylenimine (PEI) was added gradually to a final concentration of 0.25% (w/v) at 4°C while stirring to precipitate the contaminating nucleic acids. The PEI-containing suspension was clarified by centrifugation at 75,000 x g at 4°C for 30 min. Ammonium sulfate was gradually added to 60% saturation at 4°C while stirring to precipitate the protein and get rid of the PEI. The solution was centrifuged at 75,000 x g at 4°C for 30 min to collect the protein pellet. The pellet was resuspended in Buffer B [20 mM HEPES, 20% glycerol, 1 mM BME, 500 mM NaCl, at pH 7.5] containing 5 mM imidazole. Nickel-nitrilotriacetic (Ni-NTA) resin (Thermo Fisher Scientific Inc.) was equilibrated with 10 column volumes (c.v.) of Buffer B containing 5 mM imidazole at 1 mL/min using a peristaltic pump. The protein load was passed through the equilibrated Ni-NTA resin at 1 mL/min. To remove contaminants, the loaded resin was washed with 50 c.v. and 4 c.v. of Buffer B containing 5 mM and 50 mM imidazole, respectively. Finally, 3C was eluted into multiple fractions with Buffer B containing 500 mM imidazole. The purity of the 3C fractions was evaluated by Coomassie-stained SDS-PAGE (12.5%) gel. Pure and concentrated fractions were pooled and dialyzed against Buffer C [20 mM HEPES, 20% glycerol, 1 mM BME, 1 mM EDTA, 250 mM NaCl, at pH 7.5] overnight at 4°C by using a 6-8,000 MWCO dialysis membrane (Spectrum Laboratories). Dialyzed 3C sample was centrifuged at 75,000 x g at 4°C for 30 min to remove any aggregates from the solution. The protein concentration was determined by measuring the absorbance at 280 nm (*ε_max_* = 0.008945 μM^-1^·cm^-1^) with NanoDrop 1000 (Thermo Fisher Scientific Inc.). The ionic strength of the dialyzed protein solution was confirmed with a conductivity meter. Agarose gel electrophoresis was run to check for any contaminating nucleic acids. To assess the homogeneity of the protein sample, dynamic light scattering (DLS) experiments of the purified 3C (2.0 mg/mL) were performed with Viscotek 802 DLS (Malvern Instruments Ltd.) at 20°C. Flash frozen aliquots were stored at −80°C until ready to use. The ProtParam tool (ExPASy Bioinformatics Resource Portal) was used to calculate the physical/chemical parameters of the protein.

#### Expression and purification of PLCδ1 PH domain

The pSUMO-PLCδ1 PH expression plasmid was transformed into Rosetta(DE3) competent cells. The rest of the expression details are identical to that of 3C. The cell pellet was lysed and the His6-SUMO-PLCδ1 PH domain was purified via nickel affinity chromatography as described for 3C. Note that BME concentration was increased to 2 mM for each step to avoid oxidation of the cysteine residues within the PLCδ1 PH domain. During the overnight dialysis at 4°C, ULP1 (SUMO protease) was added at 1 µg ULP1 per 1 mg protein to cleave off the His6-SUMO from the PLCδ1 PH domain containing an authentic N-terminus. We observed that PLCδ1 PH domain itself binds to the nickel resin, accordingly we were unable to use a 2^nd^ nickel column to remove the His6-SUMO. Taking advantage of the differences in isoelectric points of PLCδ1 PH domain (pI = 8.4) and the His6-SUMO (pI = 5.7), we used Q-Sepharose (anion exchange) resin to separate the two. After the dialysis and the cleavage were complete, the resulting solution was diluted 10-fold with Buffer D [20 mM HEPES, 20% glycerol, 2 mM BME, 1 mM EDTA, at pH 7.5] to bring down the NaCl concentration to 50 mM. Q-Sepharose resin, at a bed volume assuming 30 mg/mL binding capacity, was equilibrated with 10 c.v. of Buffer D containing 50 mM NaCl. The protein solution was passed through the equilibrated Q-Sepharose resin at 1 mL/min. The flow-through contained the PLCδ1 PH domain. The resulting flow-through was concentrated ~30-fold with Vivaspin-20 (3,000 MWCO) centrifugal concentrators (Sartorius Stedium Biotech). The concentrated protein was re-dialyzed in Buffer D containing 250 mM NaCl. The protein concentration was determined by measuring the absorbance at 280 nm (*ε_max_* = 0.02349 μM^-1^·cm^-1^) with NanoDrop 1000. Homogeneity and purity of the PLCδ1 PH domain were assessed same as in the case of 3C.

#### Isotopic labeling and sample preparation for NMR

^15^N-labeled 3C expression was performed in autoinducible minimal media as previously described (Studier, 2005). Purification was performed as described above. Purified 3C fractions were concentrated using Vivaspin-20 centrifugal concentrators (Sartorius Stedium Biotech) and buffer exchange was performed using Zeba spin desalting columns (7,000 MWCO) (Thermo Scientific). Experimental buffer conditions were 10 mM HEPES pH 8.0, 50 mM NaCl, 10% D_2_O. Final concentrations were about 0.2 mM, 2.0 mM and 0.2 mM for 3C protein, dibutyl-PI4P, and 11-mer RNA (5′-AGU UCA AGA GC-3′), respectively.

#### NMR Spectroscopy

Resonance assignments were obtained from BioMagResBank database (BMRB ID 15222) (Amero et al., 2008). ^1^H-^15^N HSQC NMR spectra were recorded at 25°C on a 600 MHz Bruker Avance III spectrometer equipped with a 5mm "inverse detection" triple resonance (^1^H/^13^C/^15^N) single axis gradient TCI cryoprobe. NMRPipe and NMRView software were used to process and analyze the NMR spectra (Delaglio et al., 1995; Johnson and Blevins, 1994). Chemical shift perturbations were calculated using the equation below.

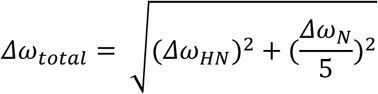

where Δω_total_ is the overall change in chemical shift, Δω_HN_ is the change in chemical shift in the amide proton dimension and Δω_N_ is the change in chemical shift in the nitrogen dimension. Chemical shift perturbations at least one standard deviation above average were considered to be substantial (Δω_total_ = 0.019 for 3C:PI4P-diC_4_ binding).

#### Fluorescence polarization-based phosphoinositide binding assay

The measurements were performed as previously described (Kolli et al., 2015). To test PIP-binding specificity, 5 µM of 3C or its mutants were added into a solution containing 0.4 nM of BODIPY-FL-labeled PIPs in a binding buffer [20 mM HEPES and 10 mM NaCl, at pH 7.5] at a 100 μL final reaction volume. Due to tight binding, PLCδ1 PH domain PIP-specificity was assessed at 100 mM NaCl and in the presence of 34 nM protein. For competition experiments, unlabeled dibutyl PI (Echelon-Inc. P-0004) and PI(4,5)P_2_ (Echelon Inc. P-4504) were titrated into a solution containing a pre-formed 3C-RNA complex. 1 μM of 3C and 0.4 nM of 3′-fluorescein (FL) labeled 11-mer RNA (5′-AGU UCA AGA GC-3′-FL), corresponding to the PV oriI sequence, were added to the binding buffer. Unlabeled dibutyl-PI(4,5)P_2_ was added last and then the milli-polarization value was recorded. Data from protein titration experiments was fit to a hyperbola. Graphs were generated by GraphPad Prism v.6 software. Error bars represent the SEM.

#### Small unilamellar vesicle (SUV) preparation

Lipids were mixed at the desired mole ratio in chloroform in a glass scintillation vial. The chloroform was removed by continuously purging the vial with N_2_ gas. Desiccation was performed under vacuum for 2-3 hours to remove any residual organic solvent. The dried lipid films were hydrated with 20 mM HEPES, 100 mM NaCl, at pH 7.0 followed by sonication in a water bath at room temperature to obtain 0.5 mg/mL lipid suspensions. These suspensions were then subjected to 10 freeze−thaw cycles with liquid N_2_ and 40°C water bath, and 10 extrusion cycles through a 100 nm track-etched polycarbonate membranes (Whatman) using a LIPEX extruder (Northern Lipids Inc.). The size and homogeneity of the SUVs were determined via Viscotek 802 DLS. To assess the quality, fluidity, and the ability of SUVs to fuse on a glass substrate, fluorescence recovery after photobleaching (FRAP) experiments were conducted. The diffusion constant for all of the vesicles prepared was within an acceptable range (>1.0 µm^2^/sec). In order to determine the pKa of the oSRB-POPE fluorescent probe within the SLBs in the presence or absence of phosphoinositides, pH titration experiments were conducted. This allowed us to make sure that the presence of phosphoinositides does not significantly change the dynamic range of the fluorescent probe.

#### Glass cleaning procedure

The glass substrates used for supporting fluid lipid bilayers were first boiled in a 7-fold diluted 7X cleaning solution (MP Biomedicals) and water for one hour. This was followed by rinsing the glasses with copious amounts of purified water before drying thoroughly with nitrogen gas. The coverslips were then annealed for five hours at 550°C before being stored until use.

#### Supported lipid bilayer (SLB) binding experiments

A microfluidic platform was employed to test the interaction between 3C and PIP-containing SLBs. PDMS cover block served as the ceiling of the device. The fabrication of the PDMS device was described previously (Jung et al., 2009). SLBs were formed on the glass floor of each microchannel by spontaneous fusion of the SUVs. Running Buffer [20 mM HEPES, 100 mM NaCl, at pH 7.0] was flowed through each channel for 30 min to get rid of excess vesicles. In order to equilibrate the SLBs to the experimental condition, Running Buffer containing 5 mM magnesium acetate was flowed through each channel for 30 min. The fluorescence intensity obtained post-equilibration step served as a reference for each channel. 3C dilutions were prepared using the Running Buffer containing 5 mM magnesium acetate as a diluent. SLBs containing 99.5 mol% POPC and 0.5 mol% *o*SRB-POPE, the pH-sensitive fluorescent lipid, served as a negative control. 3C was flowed until the fluorescence intensity in each channel stabilized (30-45 min). Change in fluorescence intensity, normalized to the reference (no 3C) channel, was plotted as a function of 3C concentration and then fit to a Langmuir isotherm using the equation below.

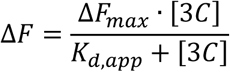

where *F_max_* represents the normalized fluorescence intensity value at saturation level and *K_d,app_* represents the apparent dissociation constant. Graphs were generated by GraphPad Prism v.6 software. Error bars represent the SEM.

Images were taken with an Axiovert 200M epifluorescence microscope (Carl Zeiss Microscopy) equipped with an AxioCam MRm camera (Carl Zeiss Microscopy) and X-Cite 120 (Excelitas Technologies Corp.) light source, was used to take fluorescence images. A 10X air objective was used for imaging along with Alexa 568 filter set (Carl Zeiss Microscopy) with an excitation and emission at 576 nm and 603 nm, respectively. Exposure times (200 msec/exposure) and the number of exposures were kept to an absolute minimum to avoid photobleaching. AxioVision LE64 v.4.9.1.0 software (Carl Zeiss Microscopy) was used to process the images.

#### Photomask design

The microfluidic device was designed with a drafting software (AutoCAD v.2016). This device consists of eight channels with independent inlets and outlets. Each channel has a dimension of 1 cm x 50 µm (length x width) and each channel is separated from each other by a 25 µm gap. The design with black background and clear features was printed at a 20,000 dpi resolution on a transparent mask (5 x 7 in) by CAD/Art Services.

#### Silicon mold fabrication

The mold containing the microfluidic patterns was fabricated in the Nanofabrication Laboratory at Penn State in University Park, PA. A 4-inch silicon wafer was dehydrated (1 min at 95°C). The SU8-50 (MicroChem Corp.), negative tone photoresist, was poured onto the center of the wafer via static dispensing method, spun (5 sec at 500 rpm, 35 sec at 3,000 rpm), pre-baked (5 min at 65°C), soft-baked (15 min at 95°C), exposed to UV light with the MA/BA6 mask aligner for 1 min (4 x 15 sec/exposure) at a power density of 8.0 mW/cm^2^ to produce a positive relief of photoresist on the wafer, and then post-baked (1 min at 65°C and 4 min at 95°C) to selectively cross-link the UV-exposed portions of the film. The wafer was developed in an SU8 developer for 6 min (without agitation), rinsed with isopropyl alcohol (IPA), and dried with N_2_ gas. The wafer with photoresist pattern was hard-baked for 30 sec at 65°C, 30 sec at 95°C, and 1 min at 150°C.

#### Molecular Dynamics Simulations of PV 3C-membrance interaction

In the simulations, a membrane consisting of 244 POPC and 20 PI4P lipid molecules, and a membrane composed of 244 POPC and 20 PI(4,5)P_2_ were prepared. The model membranes were created by the CHARMM-GUI (Wu et al., 2014) and equilibrated for 50 ns. The protein was placed above the membrane with the center of mass of the protein about 5 nm away from the center of membrane in the Z direction. The closest distance between the protein and the lipid molecules is around 0.5 nm. The starting orientation of the protein towards the membrane is based on the docking prediction, with few vital PIP-interacting residues such as R13 and R84 facing the membrane. The protein and membrane were solvated in a box of TIP3P water. Sodium and chloride ions were added at a near-physiological ion concentration of 150 mM.

The CHARMM36 force field was used for the simulation atoms (Huang and MacKerell, 2013; Klauda et al., 2010). A time step of 2 fs was employed. The van der Waals interactions was cut at 1.2 nm and the electrostatic interactions was treated with Particle-Mesh Ewald (PME) method. The SHAKE algorithm was used for length constraint on bonds involving hydrogen. The simulations were carried out under conditions of constant temperature at 310 K and constant pressure at 1 bar. The simulation system was subjected to energy minimization for 2000 steps, followed by 1 ns constraint simulation with a harmonic potential applied on protein atoms. The production simulations of 400 ns in length were then performed. The initial 30 ns of all simulations was simulated using the NAMD/2.10 package (Phillips et al., 2005). Then it was moved to the Anton supercomputer that is optimized for MD simulation for another 370 ns simulation (Shaw et al., 2009). The trajectories of the last 300 ns were used for analysis.

### Quantification and Statistical Analysis

Statistical analysis and nonlinear regression was provided by GraphPad Prism v.6. Error bars represent the SEM.

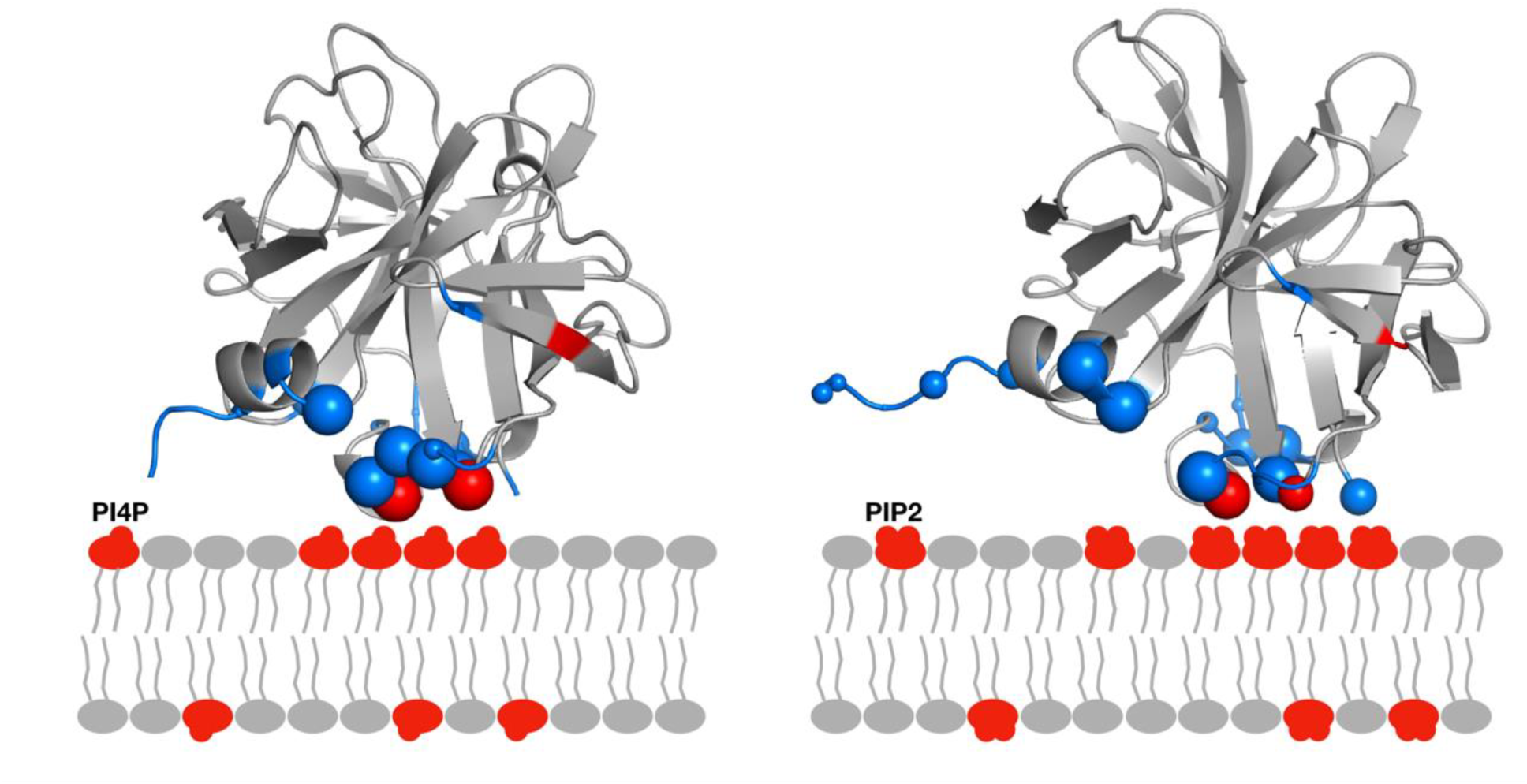
Graphical Abstract.

